# Unravelling individual rhythmic abilities using machine learning

**DOI:** 10.1101/2023.03.25.533209

**Authors:** Simone Dalla Bella, Stefan Janaqi, Charles-Etienne Benoit, Nicolas Farrugia, Valentin Bégel, Laura Verga, Eleanor E. Harding, Sonja A. Kotz

## Abstract

Humans can easily extract the rhythm of a complex sound, like music, and move to its regular beat, for example in dance. These abilities are modulated by musical training and vary significantly in untrained individuals. The causes of this variability are multidimensional and typically hard to grasp with single tasks. To date we lack a comprehensive model capturing the rhythmic fingerprints of both musicians and non-musicians. Here we harnessed machine learning to extract a parsimonious model of rhythmic abilities, based on the behavioral testing (with perceptual and motor tasks) of individuals with and without formal musical training (*n* = 79). We demonstrate that the variability of rhythmic abilities, and their link with formal and informal music experience, can be successfully captured by profiles including a minimal set of behavioral measures. These profiles can shed light on individual variability in healthy and clinical populations, and provide guidelines for personalizing rhythm-based interventions.

## Introduction

When speaking, playing a musical instrument, or walking in nature, we naturally coordinate our actions with what we perceive. Music is an excellent model for studying the link between perception and action. Listening to music often urges us to move. Sometimes we can choose to deliberately align our movements to the beat of music like we do in dance. How can we explain this widespread tendency to move to music? Musical features such as its regular temporal structure (rhythmic complexity, syncopation), but also its pitch structure (harmonic complexity) are particularly conducive to movement (Matthews et al., 2019; Witek et al., 2014). An explanation of this tight link between musical rhythm and movement can also lie in the structure and functioning of our brains (Chen et al., 2008; Grahn & Brett, 2007; Janata et al., 2012; Zatorre et al., 2007). Regions of the brain typically underpinning motor control, such as the basal ganglia and motor cortical areas, are surprisingly engaged when we mere listen to a rhythmic sequence in the absence of motor movement (Cannon & Patel, 2021; Chen et al., 2008; Grahn & Brett, 2007; Kotz et al., 2018; Nozaradan et al., 2017).

Humans – musicians and non-musicians alike – are well equipped to extract the regular pulse of music (i.e., the beat) from a complex auditory sequence and align their movements to this pulse by foot tapping, dancing, or walking (beat perception and synchronization; BPS; Damm et al., 2020; Patel & Iversen, 2014; Tranchant et al., 2016). The majority in the general population can track the beat of music and move along with it (Sowiński & Dalla Bella, 2013; Tranchant et al., 2016). Matching movements to the beat is possible because the temporal dynamics of rhythm drives internal neurocognitive self-sustained oscillations underpinning beat perception (Fujioka et al., 2012; Large & Jones, 1999; Nozaradan et al., 2011). This underlying process, called entrainment, generates temporal expectations which influence motor control, by allowing the alignment of movements to the anticipated beat times.

BPS abilities are thought to be universal skills that can be refined by musical training. Musicians outperform non-musicians in several BPS tasks. Musical training is found to improve the ability to extract the beat from a musical sequence and parse its metrical structure, and to reproduce rhythms (Chen et al., 2008; Drake & Botte, 1993; Grahn & Rowe, 2009; Kincaid et al., 2002; Nave-Blodgett et al., 2021; Smith, 1983). In addition, musicians display more precise and accurate motor synchronization to the beat than non-musicians, as shown in finger tapping to rhythmic sequences like a metronome or music (Aschersleben, 2002; Baer et al., 2013; Franĕk et al., 1991; Repp, 2010; Repp & Doggett, 2007). These effects of musical training typically manifest when participants are tested with isolated tests or a limited set of tasks, often varying from one experiment to the other. This approach, albeit valuable, may provide an incomplete picture of differences linked to musical training and a simplified characterization of individual differences. Rhythmic abilities reveal a complex structure involving different dimensions (e.g., beat-based vs. memory-based processes; Bonacina et al., 2019; Bouwer et al., 2020; Fiveash et al., 2022; Kasdan et al., 2022; A. Tierney & Kraus, 2015), compatible with the idea that there may exist multiple rhythm intelligences (Kraus, 2021). This complexity and rich structure of rhythmic abilities might not easily be captured with a few isolated tasks. There is a need of a more general, and at the same time parsimonious, way to account for individual differences in rhythmic abilities. This approach would call for multiple testing, and modelling of both the differences resulting from musical training and the fluctuations of BPS capacities in the general population.

Even in the absence of musical training, individuals can differ significantly in BPS abilities (Sowiński & Dalla Bella, 2013; Tranchant et al., 2016). A few single-case studies on beat-deaf individuals or poor synchronizers (Bégel et al., 2017; Palmer et al., 2014; Phillips-Silver et al., 2011; Sowiński & Dalla Bella, 2013), using motor tasks (finger tapping; Repp, 2005) and/or beat perception tests (e.g., the Beat Alignment Test; Iversen & Patel, 2008), reveal that either beat perception, synchronization, or both can be selectively impaired in healthy individuals. Rhythm perception and production are often both impaired in beat-deaf individuals (Palmer et al., 2014; Phillips-Silver et al., 2011). However, synchronization to the beat can be selectively impaired in the presence of spared beat perception (Sowiński & Dalla Bella, 2013). The reverse – poor perception with unimpaired synchronization to the beat – is also observed (Bégel et al., 2017). These individual differences are further exacerbated by disease, such as neurodegenerative and neurodevelopmental disorders (e.g., Parkinson, Benoit et al., 2014; Grahn & Brett, 2009; ADHD, Puyjarinet et al., 2017; speech and language impairments, Bégel et al., 2022; Corriveau & Goswami, 2009; Lense et al., 2021). Single-case evidence and studies of BPS in patient populations reveal intriguing dissociations between perception and production, and between beat-based and memory-based processes (see also Fiveash et al., 2022, for a recent study on a larger sample of non-musicians). These differences may translate into profiles characterizing individual differences in the general population, and potentially markers of developing or progressing impairment. However, owing to the complexity of these profiles, the task of identifying them and pinpointing the underlying mechanisms based on a few single cases is daunting. Even though single-case evidence is informative and suggestive, its generalization is not warranted. A systematic investigation in larger cohorts, relative to the performance of individuals with musical training, is still lacking.

In sum, previous studies show general differences in BPS abilities linked to musical training and mostly quantified by a limited set of tests. Evidence from several single-case studies hints at different rhythmic profiles. However, a general approach to individual differences in rhythmic abilities is still missing, which leaves several questions unanswered. Which tasks or measures of BPS are the most sensitive to differences due to musical training? Can we characterize an individual in terms of a given rhythmic profile (e.g., a signature of rhythmic abilities)? What is the weight of perceptual and motor processes in defining these profiles?

In the present study we aimed at characterizing rhythmic abilities in musicians and non-musicians using a data science approach, namely by exploiting predictive modelling with machine learning. The outcome of this approach is to define profiles of BPS abilities in musicians and non-musicians, and distill out a parsimonious model capable of accounting for individual differences in both groups. Given the multidimensionality of rhythmic abilities and to circumvent the limitations of previous studies, we submitted musicians and non-musicians to a battery of tests assessing both perceptual and motor timing abilities (Battery for the Assessment of Auditory Sensorimotor and Timing Abilities, BAASTA; Dalla Bella et al., 2017). The battery has been used extensively in the past and has proven to sensitively detect individual differences in healthy and patient populations (e.g., Bégel et al., 2017, 2022; Benoit et al., 2014; Puyjarinet et al., 2022; Verga et al., 2021). As multiple tests provide a wealth of information, we used dimensionality reduction techniques to pinpoint a minimal set of tasks and measures that allow capturing variability in BPS abilities linked to musical training. Machine learning (Hastie et al., 2009; Jones, 2014) and graph theory served to operationalize individual differences in rhythmic abilities, based on measures of BPS. We expected that both perceptual and motor measures would contribute to classify musicians and non-musicians, and that musicians would display a stronger link between perceptual and motor measures. This hypothesis is grounded in both behavioral and brain imaging evidence of a tighter coupling between perceptual and motor systems as a result of extensive musical practice (Herholz & Zatorre, 2012; van Vugt & Tillmann, 2014; Zatorre et al., 2007). Musicians tend to activate jointly auditory and motor regions when listening to sound (Chen et al., 2008; Lahav et al., 2005), or merely moving their hands (Lotze et al., 1999, 2003). An increased sensorimotor association can be found after relatively little training while learning a musical instrument (Lega et al., 2016; Wollman et al., 2018), a process which is likely involving the dorsolateral premotor cortex (Chen et al., 2012). A second goal was to examine whether non-musicians are homogenous in terms of their BPS abilities, namely if they lie on a continuum of rhythmic abilities, or rather cluster in groups. Individual differences found in single-case studies point towards different profiles, which may characterize clusters of individuals in the general population. To test this possibility, we used unsupervised learning methods (clustering).

## Results

### Defining profiles of rhythmic abilities in musicians and non-musicians

To define a signature of rhythmic abilities based on the results of BAASTA tasks that should allow teasing apart musicians from non-musicians, we processed data from 55 measures using Sparse Learning and Filtering algorithms (SLF; Combettes & Pesquet, 2007; Jiu et al., 2021). SLF aims at selecting a minimal set of variables obtained from perceptual and motor tasks of BAASTA (Perc: perceptual; M: motor) that capture the most relevant differences between musicians and non-musicians. This set of measures was then entered in a model, aimed at classifying individuals as musicians or non-musicians based on their rhythmic performance. The outcome model, including a limited set of measures, is parsimonious and affords a significant gain in statistical power relative to a model that includes all the variables. We systematically tested three classification models, by taking as input 1) the entire set of perceptual measures (7 out of 55; *Model Perc*), 2) the entire set of motor measures (48 out of 55; *Model Motor*), and 3) the selected measures resulting from Models Perc and Motor (*Model PMI*). Model PMI also included interactions between motor and perceptual measures to test the hypothesis that coupling of motor and perceptual processes is more relevant in defining the rhythmic profile of musicians than of the one of non-musicians. The procedure and the selected variables are illustrated in Figure 1. Model Perc significantly accounted for 50% of the variance (*F*(73) = 14.4, *p* < .0001), model Motor accounted for 84% of the variance (*F*(74) = 97.1, *p* < .0001), and finally model PMI accounted for 92% of the variance (*F*(70) = 99.5, *p* < .0001), relative to the prediction of a model using the entire set of variables. Accuracy for the three models is shown in Figure 1. Model PMI included a) two perceptual measures, reflecting the ability to detect whether a metronome is aligned to the beat of music or not (P1), or whether there is an irregularity in an isochronous sequence (P2), b) two motor measures, indicating the alignment of participants’ taps with the beat of a metronome (M1) and music (M2), and c) the four interactions between these measures.

**Figure 1.**
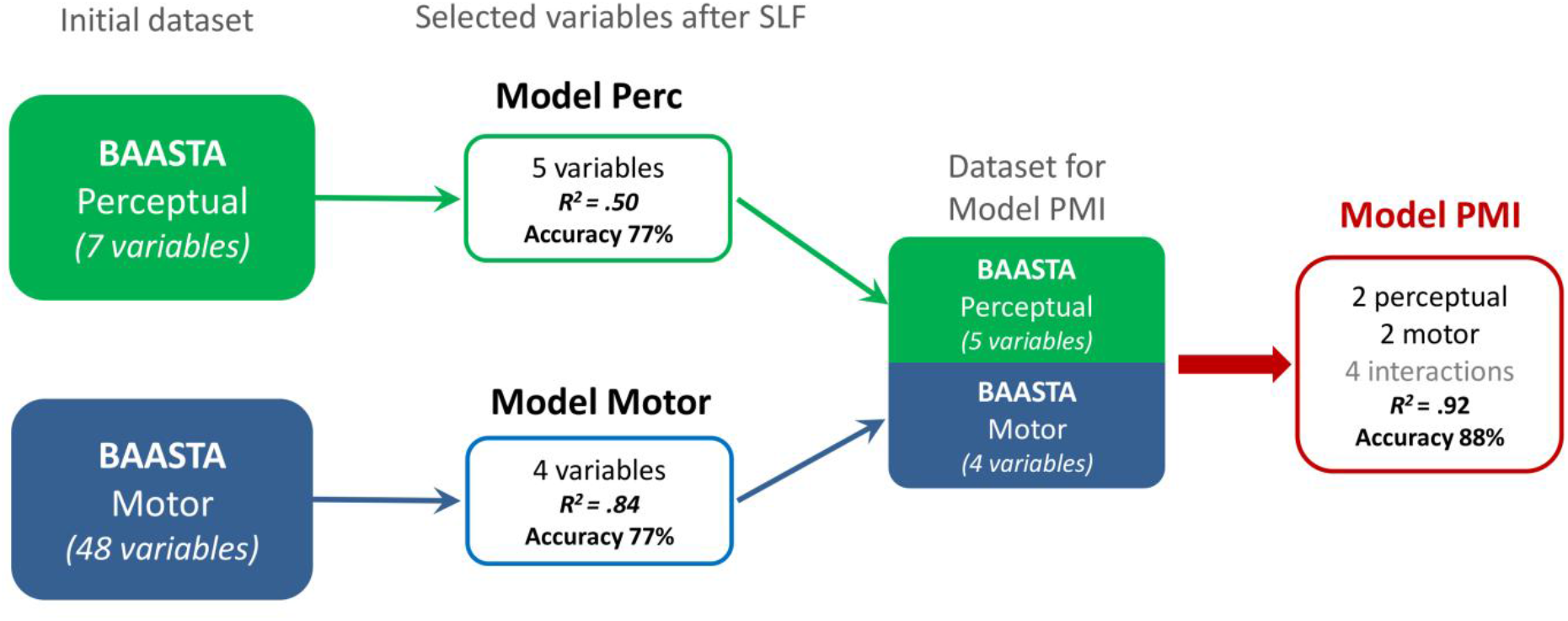
Schema of the analysis pipeline using Sparse Learning and Filtering (SLF). Selected variables for *Model Perc*: 3 variables from the Beat Alignment Test (BAT_slow_Dprime, BAT_med_Dprime, BAT_fast_Dprime), 1 from Duration discrimination (Dur_discrim_Thresh), and 1 from Anisochrony detection with music (Anisoc_det_music_Thresh). *Model Motor*: 2 variables from Paced tapping with tones (Paced_metro_750_Vector_dir, Paced_metro_750_Vector_len), 1 from Paced tapping with music (Paced_music_ross_Vector_dir), and 1 from Unpaced tapping (Unpaced_slow_CV). *Model PMI*: 2 variables from perceptual tests (*P*_1_ – BAT_slow-Dprime; *P*_2_ – Anisoc_det_music_Thresh), 2 from motor tests (*M*_1_ – Paced_metro_750_Vector_dir; *M*_2_ – Paced_music_ross_Vector_dir), and the 4 interactions *P*_1_\**M*_1_, *P*_1_\**M*_2_, *P*_2_\**M*_1_, *P*_2_\**M*_2_. Accuracy is calculated based on the Test phase of the classifier validation. Full description of the variables, and confusion matrices for the three models at Train (60% of the dataset), Validation (20%) and Test (20%) phases are provided in Supplementary materials.

We tested the capacity of the three models to discriminate between musicians and non-musicians by taking the outcome of the models (value from 0 to 1, with 0 indicating non-musicians, and 1, musicians) using the entire dataset (i.e., including the three phases of the classifier validation). Model PMI was superior compared to the other two models in teasing apart musicians and non-musicians (Figure 2a; Model x Group interaction; *F*(2, 154) = 3.8, *p* < .05; model Perc (*t*(66.9) = 5.2, *p* < .0001, *d* = 1.3), model Motor (*t*(64.3) = 6.2, *p* < .0001, *d* = 1.5); model PMI (*t*(64.0) = 7.0, *p* < .0001, *d* = 1.8). Even though the distributions of the model PMI predictions were partly overlapping (Figure 2b), the model could classify musicians and non-musicians very successfully (Figure 2c). Equation 1 below allowed the classification of musicians and non-musicians (prediction = *ỹ*) for model PMI.

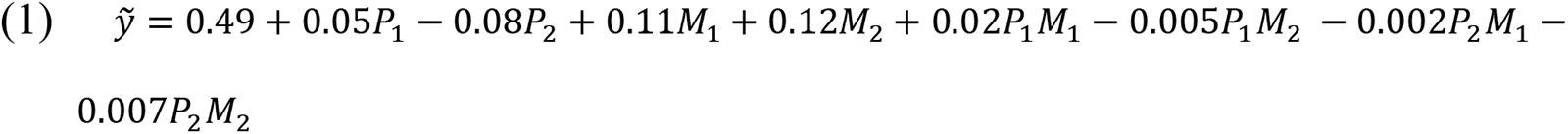

**Figure 2.**
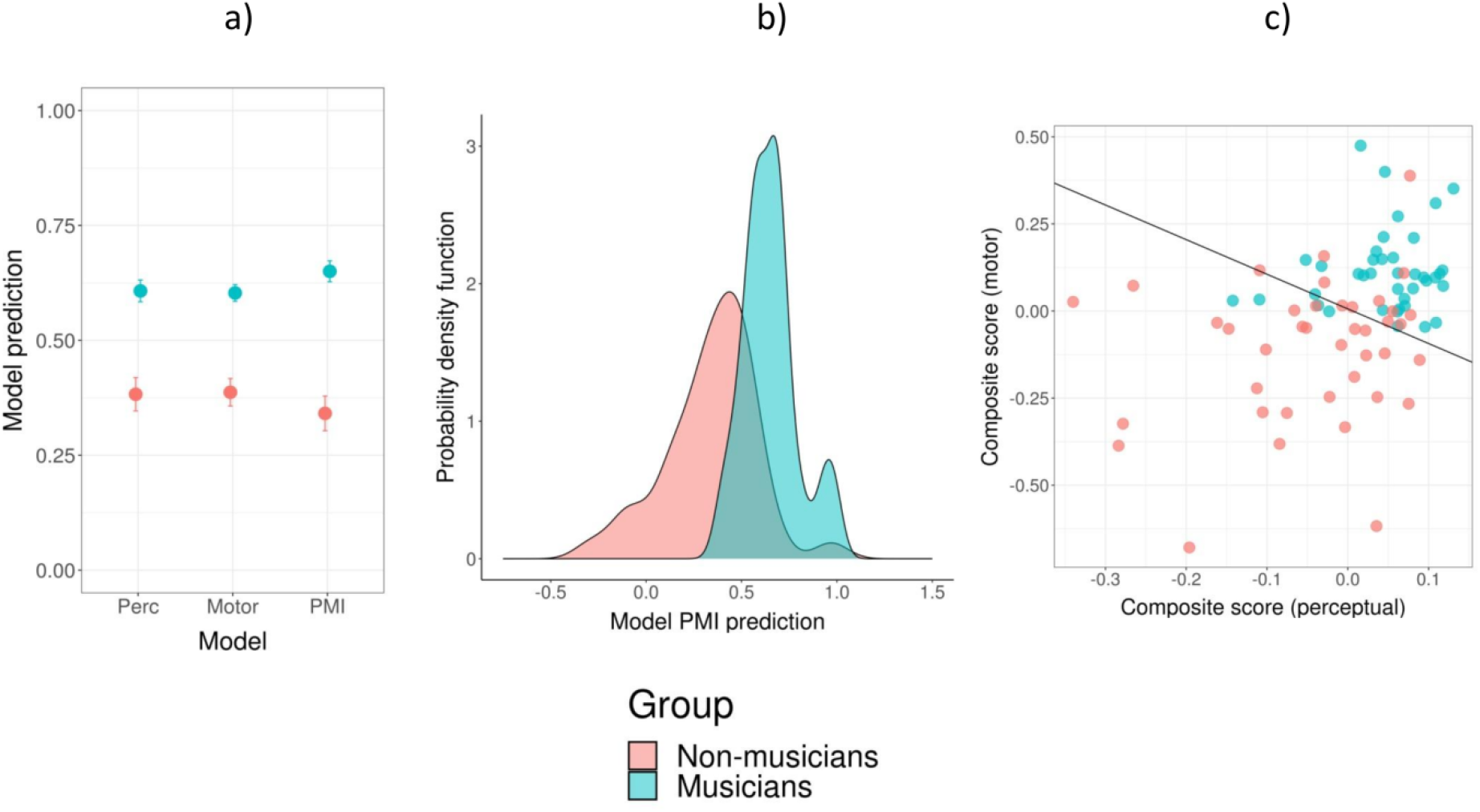
Classification performance of the three models. a) Comparison of the Perc, Motor, and PMI models predictions (0 = non-musicians; 1 = musicians). Error bars represent SEM. b) Probability density for musicians and non-musicians obtained with model PMI. c) Scatter plot showing the individual predictions based on model PMI. For simplicity, the predictions are presented as a projection of two composite scores representing the linear portions of the prediction model, referring to the perceptual measures (perceptual score = 0.05*P_1_*– 0.08*P_2_*) and the motor measures (motor score = 0.11*M_1_*+ 0.12*M_2_*), without the interactions.

Model PMI, including two perceptual variables, two motor variables and the four interactions yielded the best classification results. To gain a better understanding of the rhythmic signature of musicians and non-musicians, we examined the contribution to the classification of each variable and interaction independently for each group. We calculated the correlation between each variable and the prediction separately for musicians and non-musicians, leading to a measure of explained variance. With this method we obtained two profiles characterizing musicians and non-musicians (Figure 3a and 3b). The contribution of each component (perceptual, motor, and their interaction) to the profiles of musicians and non-musicians is illustrated in Figure 3c. The motor measures contributed to the profiles more than perceptual measures in both musicians (*χ*^2^(1) = 20.8, *p* < .0001) and non-musicians (*χ*^2^(1) = 22.4, *p* < .0001). Differences between the two profiles regarding the contribution of individual variables and interactions are apparent. Interactions between perceptual and motor variables play a more important role in the definition of musicians’ profile than for non-musicians (*χ*^2^(1) = 5.2, *p* < .05). The motor component may contribute more to the definition of non-musicians, but this difference did not reach statistical significance (*χ*^2^(1) = 3.5, *p* = .11).

**Figure 3.**
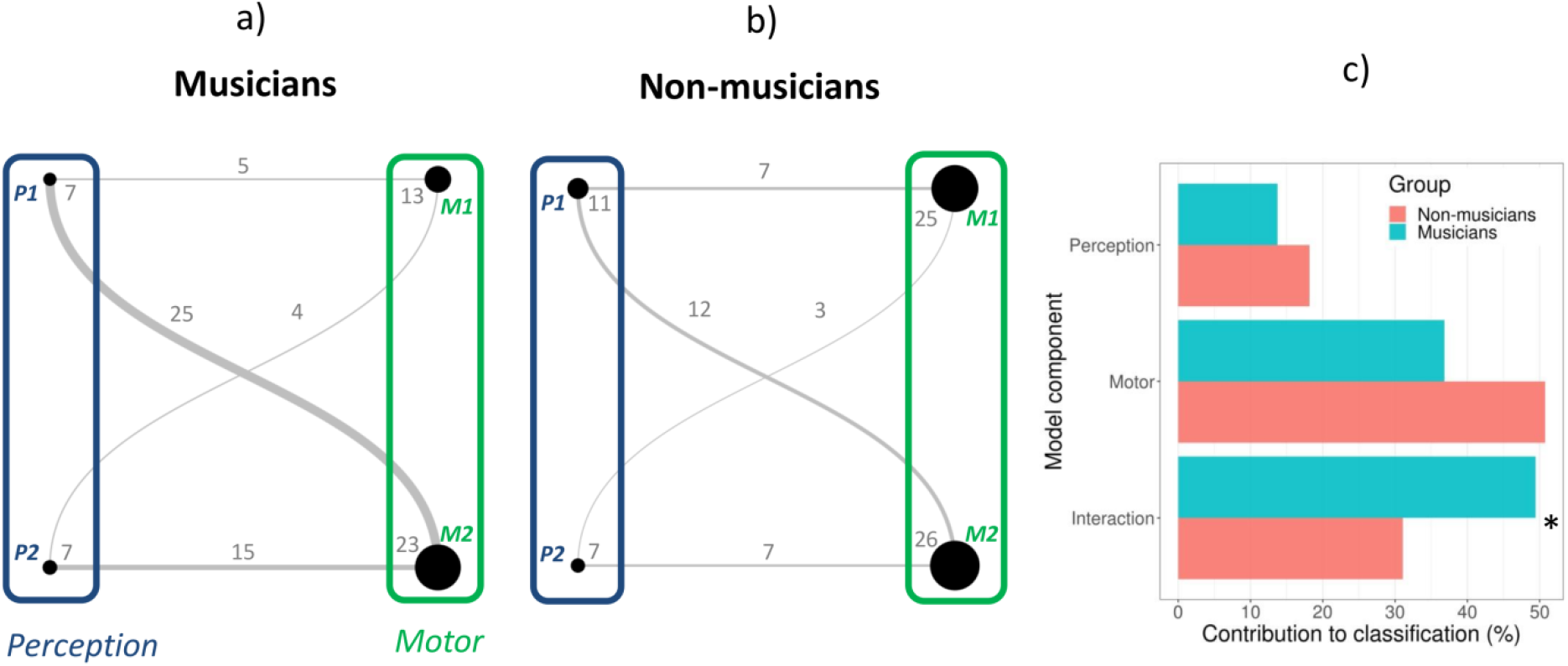
Profiles of rhythmic abilities for musicians (panel a) and non-musicians (panel b) based on model PMI, expressed as undirected graphs. The nodes’ size reflects the contribution of the variable to the definition of the group (proportion of variance) and the edges’ widths to the contribution of the interactions. c) Comparison of the contribution of the model component (perceptual, motor, interaction) to the profiles of musicians and non-musicians. P = perceptual; M = motor. Numbers refer to the specific variable (*P1* – BAT_slow-Dprime; *P2* – Anisoc_det_music_Thresh; *M1* – Paced_metro_750_Vector_dir; *M*2 – Paced_music_ross_Vector_dir). * *p* < .05.

These findings reveal that a minimal set of rhythmic measures and their interactions can successfully differentiate musicians from non-musicians, as illustrated by undirected graphs. A limitation of this representation, however, is that it does not reflect the sign of the relation, positive or negative, between the variable and the predicted group. Moreover, one might wonder whether each component of the model can explain individual variability within each group. To address this point, we examined the relation between each variable and interaction and the prediction, separately for each group (Figure 4). Each component of the profile can explain differences within each group. The only exception is the interaction between P2 and M1, which just failed to reach significance in explaining the variability for non-musicians. Notably the motor measures alone could explain large portions of intra-group individual variability (63% for non-musicians and 46% musicians). Moreover, interactions between perceptual and motor variables could capture a significant amount of individual variability in both groups. This further supports the inclusion of interactions in the PMI model. Also worth noting is the sign of the relation between the components and the prediction. For both groups, improved performance (i.e., an increase of P1, M2, or M2; a decrease of P2, indicating a lower detection threshold) was associated with greater rhythmic abilities. With interactions, different relations were found for musicians and non-musicians. For example, a decrease of the value of the interaction between P1 and one of the two motor variables was associated with greater rhythmic abilities in non-musicians; however, the opposite relation was observed in musicians. Thus, interactions between perceptual and motor measures of rhythmic abilities may contribute differently to the characterization of musicians and non-musicians.

**Figure 4.**
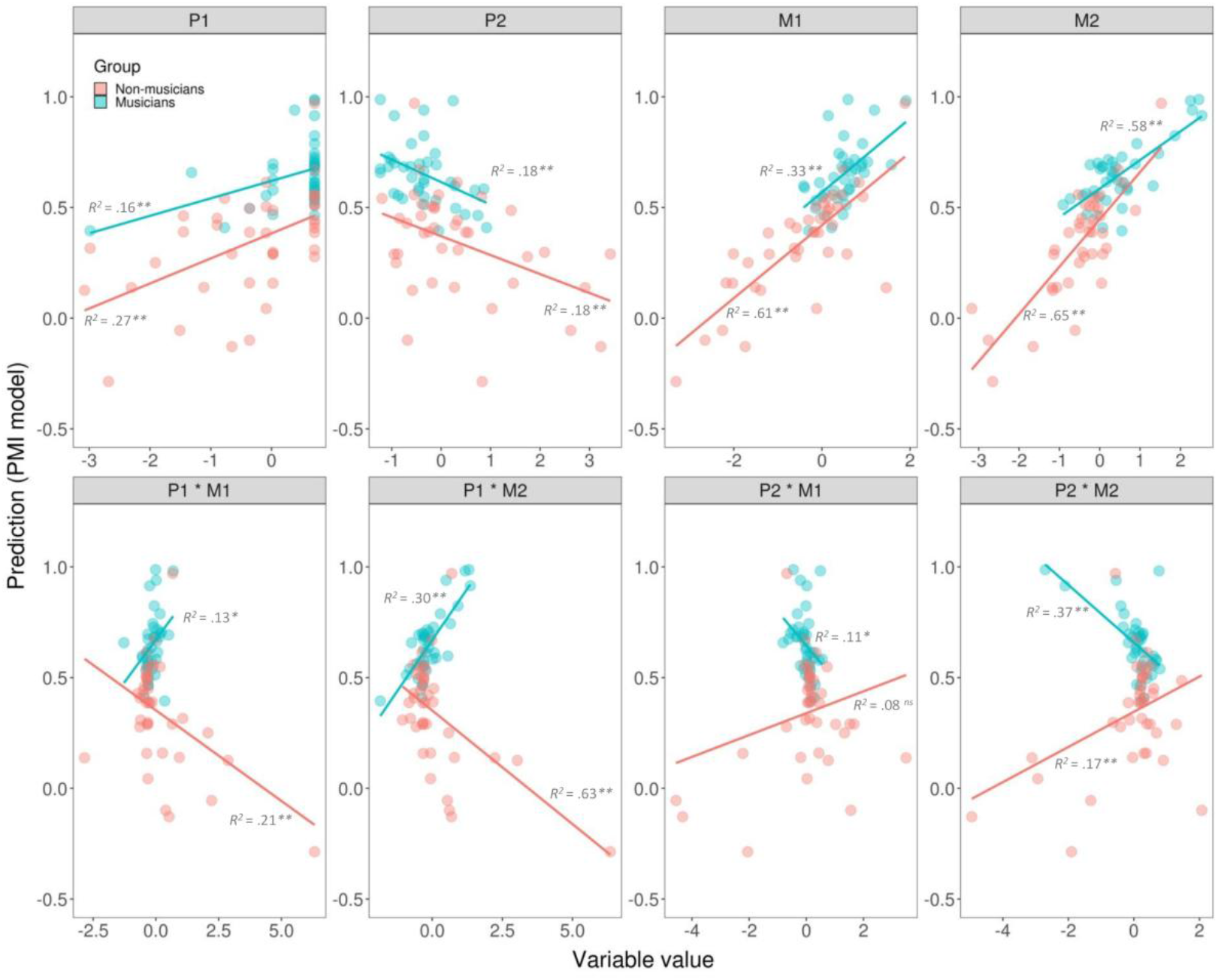
Scatter graphs showing the relation between individual elements of the profiles of rhythmic abilities (measures and their interactions) and the prediction of model PMI, separately for musicians and non-musicians. Explained variance for each group is reported. ** *p* < .01. * *p* < .05 ns = not significant.

### Profiles of rhythmic abilities for non-musicians

An inspection of measures of rhythmic abilities in non-musicians reveals considerable variability in this group. To assess whether non-musicians’ rhythmic performance was homogeneous or rather different clusters (and corresponding profiles) may emerge, we used model PMI to examine the distribution of rhythmic skills in this group. Building on graph theory, we extracted two clusters from the non-musicians group using modularity analysis (Newman, 2006; Rubinov & Sporns, 2010). This procedure led to the detection of two clusters (Subgroup 1, Subgroup 2) that maximized modularity. Subgroup 1 included 20 participants (9 females, mean age = 24.1 years, *SD* = 4.7), and Subgroup 2 was formed by 20 participants (10 females, mean age = 22.1 years, *SD* = 2.6). The two subgroups did not differ significantly in terms of years of formal musical training (mean = 0.8 years, *SD* = 1.5, for Subgroup 1; mean = 1.8 years, *SD* = 2.4, for Subgroup 2; *t*(31.8) = 1.6, *p* = .14, *d* = .5). However, Subgroup 2 showed more years of informal musical activities (mean = 1.7 years, *SD* = 2.9, for Subgroup 1; mean = 5.0 years, *SD* = 4.3, for Subgroup 2; *t*(33.3) = 2.8, *p* < .01, *d* = 1.0).

Once we uncovered the two clusters of non-musicians, we used the variables of model PMI to classify the two subgroups (see equation 2 below). This new model could classify the two subgroups with an accuracy of 100% (in the Test phase of the classifier validation; the confusion matrices at Train, Validation and Test phases are provided in Supplementary materials). The performance of the model is seen in Figure 5.

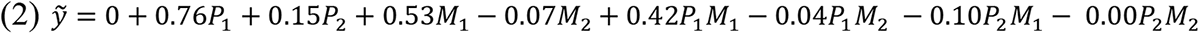

**Figure 5.**
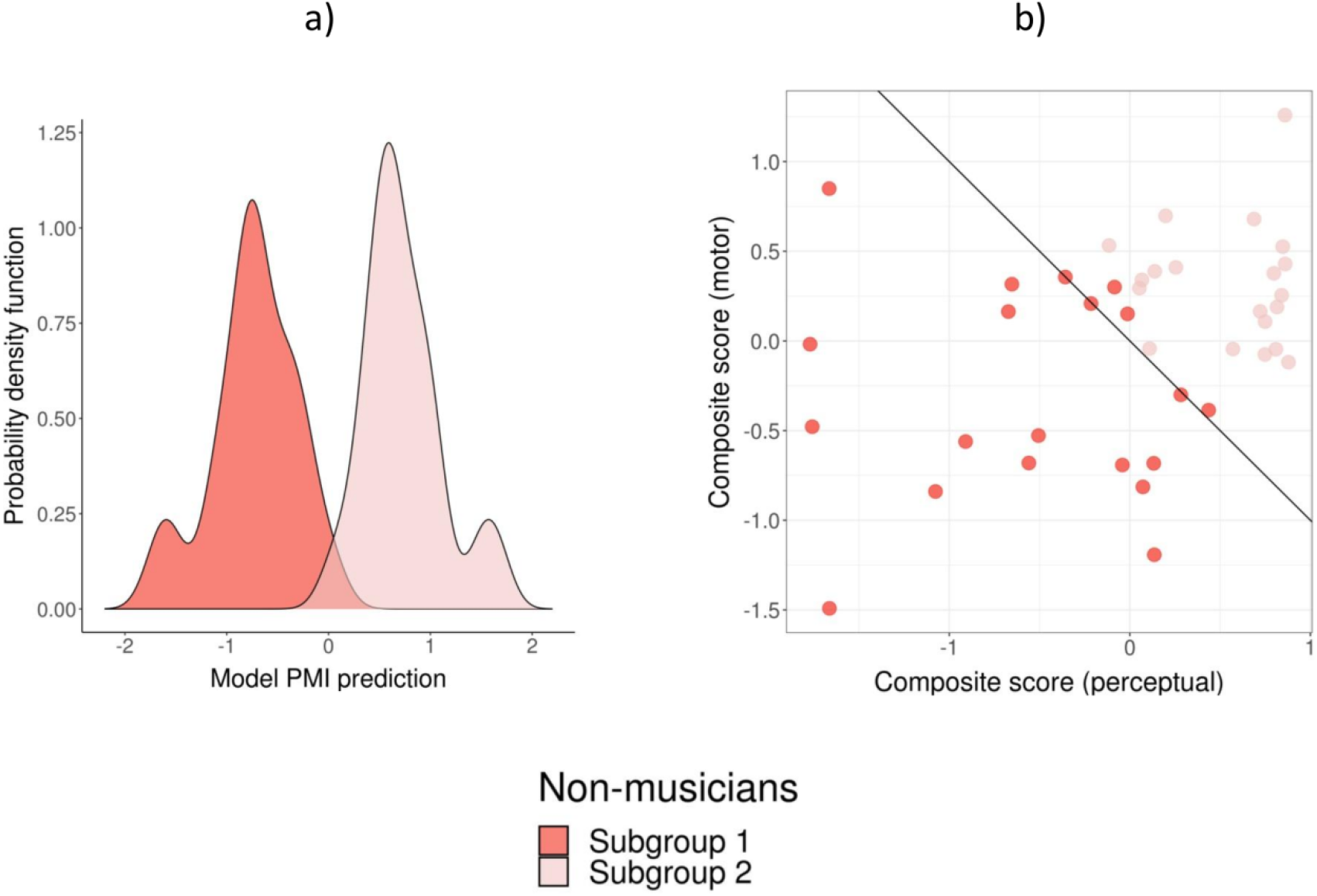
Classification performance of Model PMI for the two subgroups of non-musicians (– 1 = Subgroup 1; 1 = Subgroup 2). a) Probability density for non-musicians in Subgroup 1 and Subgroup 2. b) Scatter plot showing the individual predictions based on the model. The predictions are presented as a projection of two composite scores representing the linear portions of the prediction model, referring to the perceptual variables (perceptual score = 0.76*P_1_*+ 0.15*P_2_*) and the motor variables (motor score = + 0.53M*_1_*-0.07*M_2_*), without the interactions.

We obtained profiles for the two subgroups of non-musicians, by examining the contribution of each variable and interaction to the classification of each group (in Figure 6, panels *a* and *b*). The contribution to the profiles of Subgroups 1 and 2 of each component (perceptual, motor, and their interaction) is presented in Figure 6c. The motor measures contributed to the profiles more than perceptual measures for both Subgroup 1 (*χ*^2^(1) = 32.1, *p* < .0001) and Subgroup 2 (*χ*^2^(1) = 8.8, *p* < .01). There was a tendency toward a greater contribution of the motor component in Subgroup 1 than in Subgroup 2 (*χ*^2^(1) = 3.2, *p* = .07).

**Figure 6.**
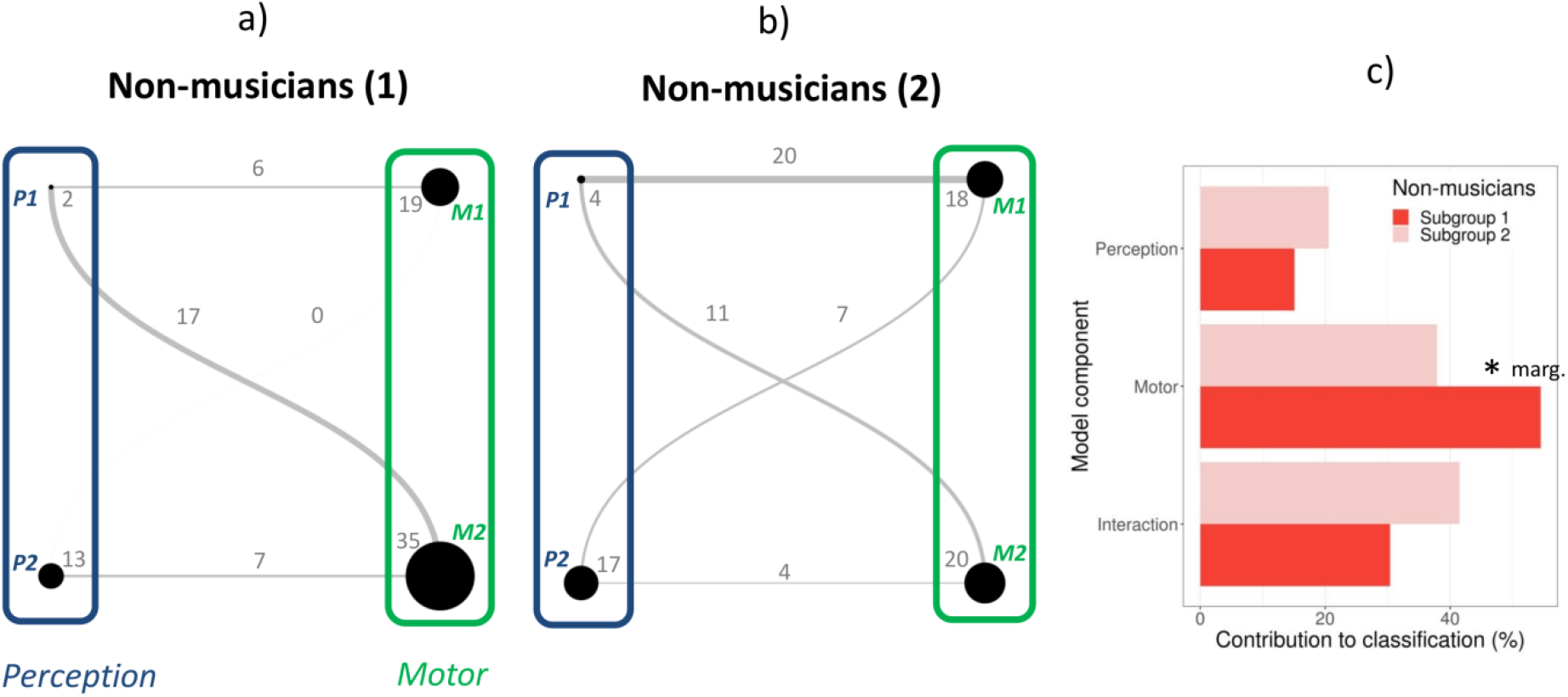
Profiles of rhythmic abilities for non-musicians in Subgroup 1 (panel a) and Subgroup 2 (panel b) expressed as undirected graphs. The nodes’ size reflects the contribution of the variable to the definition of the group (proportion of variance) and the edges the contribution of the interactions. (c) Comparison of the contribution of the model component (perceptual, motor, interaction) to the profiles of Subgroups 1 and 2). P = perceptual; M = motor. Numbers refer to the specific variable (*P1* – BAT_slow-Dprime; *P2* – Anisoc_det_music_Thresh; *M1* – Paced_metro_750_Vector_dir; *M*2 – Paced_music_ross_Vector_dir). * marg. = marginally significant, *p* = .07.

To test whether each component of the non-musicians’ subgroup profiles can explain individual variability within each group, we examined the relation between each variable and interaction and the prediction separately for each subgroup (Figure 7).

**Figure 7.**
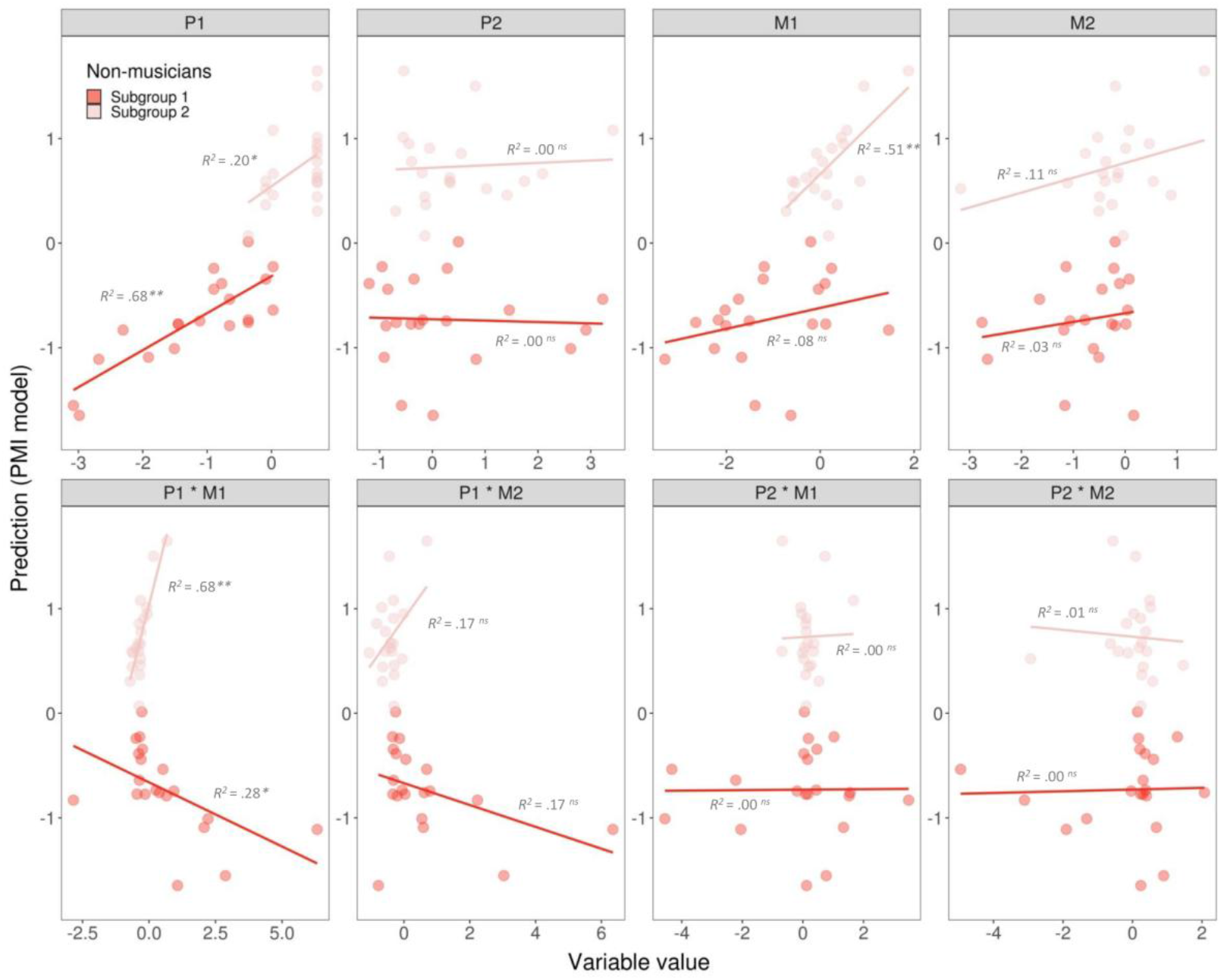
Scatter plots showing the relation between individual elements of the profiles of rhythmic abilities (variables and interactions) and the prediction of model PMI, separately for non-musicians in Subgroups 1 and 2. Explained variance for each group is reported. ** *p* < .01. * *p* < .05 ns = not significant.

Only a few components of the profiles could explain the variability in each Subgroup, in a rather specific way. Notably, the two components P1 and M1, and their interaction, were the only ones that explained the variability within the two subgroups. Variability within Subgroup 1 was strongly related with component P1, explaining 68% of its variance, and with the interaction between component P1 and M1. Variability within Subgroup 2 is mostly explained by component M1 (51% of the variance) and the interaction between M1 and P1 (68%). Finally, because the two subgroups of non-musicians differed in terms of informal musical activities, we tested whether variability in each of the two subgroups was related to this variable. Variability within Subgroup 1, indicated by the prediction of PMI model, was correlated with informal musical training (*r* = .48, p < .05). With increasing years of informal musical training, the prediction from the model for participants in Subgroup 1 became closer to 0, thus namely nearing the performance of Subgroup 2. This finding is coherent with the observation that Subgroup 2 displays significantly more informal musical experiences than Subgroup 1 (see above).

## Discussion

In the present study we used machine learning techniques to characterize BPS abilities in musicians and non-musicians. This approach served to operationalize individual differences in rhythmic abilities. Supervised learning was instrumental in selecting a minimal set of perceptual and motor measures from a battery of rhythmic tests (BAASTA; Dalla Bella et al., 2017), for the purpose of classifying musicians and non-musicians. We expected both perceptual and motor measures to contribute to the classification of musicians and non-musicians. In keeping with our hypothesis, a classification model including small sets of perceptual and motor measures (4 variables overall) and their interactions (model PMI) was superior to models of perceptual and motor measures on their own. The final model of rhythmic abilities was very accurate in classifying musicians and non-musicians (almost 90%) and allowed distilling separate profiles for the two groups. Profiles took the form of undirected graphs, where the nodes’ size reflects the contribution of the measure to the definition of the group (proportion of variance) and the edges the contribution of the interactions.

An inspection of the profiles of rhythmic abilities reveals that motor measures per se played a more important role (explaining more variance) than perceptual measures for both musicians and non-musicians. As expected, we found stronger interactions between perceptual and motor measures in the profiles of musicians than non-musicians. Interestingly, the components of the profiles, albeit limited in number, could capture successfully individual variability within each group. An improvement of rhythm perception (better detection of beat alignment or of deviations from isochrony) or production (i.e., a more positive relative phase between taps and the beat) increased the probability to be classified as musicians (with prediction scores closer to 1). The interactions between perceptual and motor measures also explained a significant amount of variance within each group. Notably, these interactions were differently linked to the performance of musicians and non-musicians in rhythmic tasks as shown by opposite slopes relative to the model prediction in the two groups. In sum, it is apparent that interactions between perceptual and motor measures play a critical role in defining a musician’s rhythmic profile. Taking interactions into account, both improves the classification of an individual as a musician or a non-musician and can specifically account for intra-group variability.

These findings obtained with purely behavioral measures are consistent with neuroimaging evidence of stronger coupling of auditory and motor systems resulting from extensive musical practice (Herholz & Zatorre, 2012; Zatorre et al., 2007). Musical training and performance, require the learning of specialized sensorimotor skills which put a high load on sensory, motor, and cognitive systems (Dalla Bella, 2016; Merrett et al., 2013). Expert performance in adults is grounded in extensive learning and practice of numerous skills over the course of a musician’s lifetime. The effects of both short-term and long-term musical training on brain plasticity have been widely documented and pertain to primary sensory and motor regions, as well as sensorimotor integration areas (for reviews, Dalla Bella, 2016; Herholz & Zatorre, 2012; Merrett et al., 2013; Strait & Kraus, 2014; Wan & Schlaug, 2010). For example, a model integrating these components, based on research on both vocal and instrumental performance, has been proposed by Zatorre and collaborators (Zatorre et al., 2007). The model predicts feedforward and feedback interactions involving motor and sensory regions, as well as areas devoted to sensorimotor transformation. Auditory-motor associations can emerge as the result of little training when learning a musical instrument (Wollman et al., 2018, after training non-musicians to play an MRI-compatible cello) and are thought to engage the dorsolateral premotor cortex (Chen et al., 2012). The exam of the neuronal underpinnings of auditory-motor integration when playing different musical instruments and singing (e.g., Segado et al., 2018) reveal overlaps, for example in dorsal premotor and supplementary motor cortices, thus pointing to shared mechanisms. A next step in future research should be to link the profiles of rhythmic abilities identified herein with behavioral measures with their brain substrates, to gain a better understanding of these individual differences.

A second goal of this study was to examine individual differences in non-musicians by using the same modelling approach that allowed identifying the profiles of rhythmic abilities. A critical question was whether we could identify specific clusters of individuals in non-musicians regarding BPS. By applying modular clustering, an unsupervised learning method, to the data extracted from non-musicians’ profiles, we could identify two clusters. Again, motor measures contributed more than perceptual measures to both profiles, with a slight tendency to contribute more to one of the two profiles (Subgroup1). The components of the profiles capturing the variability within each subgroup were confined to the performance in two tasks (detecting a misalignment between a metronome and the musical beat and finger tapping to a metronome), and their interaction. It is striking that one of these measures was sufficient to capture most of the variance in each group (the perceptual measure for Subgroup 1 and the motor measure for Subgroup 2), thus providing a very parsimonious metric of individual differences in the rhythmic domain. Generally, the interactions between perceptual and motor measures were less specific in capturing individual variability than what we observed when comparing musicians and non-musicians.

The two clusters of non-musicians differed in their informal musical activities. People belonging to the subgroup displaying more years of informal musical activities, albeit being classified as non-musicians, received less extreme prediction scores by the model (farther away from -1) than the other group. Consistently, the years of informal musical activities could also account for individual variability within one of the subgroups. This proves that model PMI can capture subtle differences in rhythmic abilities linked to informal musical activities, such as playing a musical instrument as an amateur musician would do. There is substantial evidence that formal musical training is not a pre-requisite to develop rhythmic abilities. Western listeners, musicians, and non-musicians alike, can acquire implicitly complex musical features (melody, harmony, rhythm) from mere exposure to music (Rohrmeier & Rebuschat, 2012). For example, non-musicians can develop implicit knowledge of pitch regularities characteristic of the Western tonal system (tonal knowledge; Tillmann, 2005; Tillmann et al., 2000) and of the temporal structure of event sequences (Bigand & Poulin-Charronnat, 2006; Terry et al., 2016; Tillmann et al., 2011). This implicit knowledge paves the way for non-musicians’ perception of the relations between musical events, thus creating expectations for upcoming events that in turn, influence their processing (Jones et al., 2002; Selchenkova et al., 2014). In sum, by mere exposure to music, Western non-musicians can acquire implicit knowledge of the different aspects of music prevalent in their culture, which, in turn, shape their perception (Bigand & Poulin-Charronnat, 2006) and production (Weiss & Peretz, 2022). Interestingly, informal music activities are likely to play an important role earlier in life during music acquisition (Hannon & Trainor, 2007). Formal musical lessons in childhood have been devoted much attention as a way to convey musical knowledge. Yet, more recently there is growing interest in the role of informal musical activities, for example, in the home environment, with evidence suggesting positive effects on both music and language development, auditory processing, and vocabulary and grammar acquisition (Politimou et al., 2019; Putkinen et al., 2013; Williams et al., 2015). Having a model of rhythmic abilities capturing the effects of informal musical activities is particularly appealing. This offers an opportunity to shed light on the role of rhythmic capacities during development, as informal musical activities are quite common, and ultimately to uncover BPS profiles associated with developmental disorders (e.g., Bégel et al., 2022; Ladányi et al., 2020; Lense et al., 2021; Puyjarinet et al., 2017). Whether the model and the selected variables are appropriate to reflect individual differences during development awaits further research.

## Conclusions and perspectives

The effects of musical training on beat perception and synchronization are well established. Musical training enhances rhythm perception and production (Chen et al., 2008; Drake & Botte, 1993; Nave-Blodgett et al., 2021), and is typically associated with better motor synchronization accuracy when tapping along with rhythmic sequences (Aschersleben, 2002; Baer et al., 2013; Franĕk et al., 1991; Repp, 2010; Repp & Doggett, 2007). However, when considering the majority of individuals without musical training our current knowledge of individual differences in BPS is mostly limited to single-case studies revealing intriguing dissociations between perception and production (Bégel et al., 2017; Palmer et al., 2014; Phillips-Silver et al., 2011; Sowiński & Dalla Bella, 2013), and group studies using a reduced set of tasks (e.g., Bonacina et al., 2019; Bouwer et al., 2020; Tierney & Kraus, 2015). An exception is a recent study by Fiveash and collaborators (Fiveash et al., 2022) in which we tested non-musicians with a large set of rhythm and perception tasks taken from different test batteries. The tasks tapped into different rhythmic competencies, such as rhythm production, beat-based rhythm perception, and sequence memory-based rhythm perception. Different patterns of performance across rhythm tasks and individuals indicated that beat-based and memory-based rhythmic competencies can be separated by behavioral testing (see also Bonacina et al., 2019; Bouwer et al., 2020; Tierney & Kraus, 2015), pointing to the complexity and multidimensionality of rhythmic abilities.

To date we are missing a general approach to rhythmic abilities (with a focus on BPS skills) that can account for differences linked to musical training and variability in non-musicians. Here we filled this gap by using machine learning techniques to extract a parsimonious model of rhythmic abilities from a large set of perceptual and motor tasks (BAASTA, Dalla Bella et al., 2017). Machine learning is perfectly suited for this task (Hastie et al., 2009; Jones, 2014), by allowing the reduction of complexity in a large multivariate dataset, while maintaining high classification accuracy, limiting overfitting, and increasing generalization to new data. As a result, a reduced set of explaining variables (i.e., a handful of perceptual and motor measures) forming a profile of rhythmic abilities is clearly easier to understand and interpret than a larger dataset. The model of rhythmic abilities we obtained (PMI), summarized by an undirected graph, was not only very successful in separating musicians and non-musicians but also its components could capture the variability of non-musicians’ performance. Based on behavioral measures, non-musicians clustered into two subgroups based on their BPS skills; these clusters differ regarding informal musical experience. Finally, perceptual and motor measures of rhythmic abilities selected for the profiles could account for within-group variability. In the end, using machine learning for the purpose of building a model of rhythmic abilities was a very fruitful endeavor. Machine learning is progressively becoming an important element of the toolbox for research in cognitive science and neuroscience (Aglinskas et al., 2022; Richards et al., 2019; Shen et al., 2022; Yarkoni & Westfall, 2017) with applications in music research (for a review, Agres et al., 2021) such as computational music analysis (e.g., music information retrieval and automatic music classification; (Lau & Ajoodha, 2022; Mueller et al., 2019; Stober et al., 2014), and more recently in music cognition (e.g., for emotion detection, Vempala & Russo, 2017). A similar approach based on machine learning and graph theory as used to model individual differences in rhythmic abilities could be purposefully extended to other music abilities such as pitch perception and production, or improvisation (e.g., Farrugia et al., 2021, for a single-case study).

By exploiting machine learning starting from a dataset issued from behavioral tests we demonstrated that the complexity of rhythmic abilities (e.g., Fiveash et al., 2022), or rhythm intelligences, (Kraus, 2021), at least when considering BPS skills, can be captured by a minimal set of behavioral measures and their interaction. Detecting profiles of rhythmic abilities is valuable to shed light on individual variability, its underlying mechanisms (e.g., neuronal, Tierney et al., 2017; genetic, Niarchou et al., 2022), and how they are affected by formal and informal musical experiences. For example, given the involvement of general cognitive processes in BPS tasks, an important line of enquiry is to examine the relation between profiles of rhythmic abilities and general cognitive functioning. Rhythmic abilities and cognitive functions such as cognitive flexibility, inhibition, and working memory tend to covary in both healthy individuals and in populations with disorders (Bégel et al., 2022; Puyjarinet et al., 2017; Tierney & Kraus, 2013); moreover, executive functions and rhythmic abilities are both enhanced by musical training (Bailey & Penhune, 2010; Zuk et al., 2014). Whether this correlation is underpinned by similar or partly distinct mechanisms is still unclear (for recent evidence pointing towards relatively independence in children with developmental dyslexia, see Bégel et al., 2022). A systematic study of the relation between profiles of rhythmic abilities and cognitive functions is likely to advance our understanding of the inter-dependence between rhythmic capacities and general cognitive functions. Finally, it is apparent that individual differences in BPS are exacerbated by diseases (e.g., Parkinson, Benoit et al., 2014; Grahn & Brett, 2009; ADHD, Puyjarinet et al., 2017; speech and language impairments, Bégel et al., 2022; Corriveau & Goswami, 2009; Falk et al., 2015; Ladányi et al., 2020; Lense et al., 2021). The proposed model and the profiles of rhythmic abilities may serve for identifying markers of impairment in clinical populations (Robinaugh et al., 2020). One of the advantages of this approach, owing to the model’s parsimony, is that profiles can be obtained via rapid testing and screening of participants with a limited set of tasks (the Beat Alignment Test, and finger tapping to music), that can be performed in 10-15 minutes. This corresponds to a gain in time over 80% relative to the full testing battery (lasting two hours; Dalla Bella et al. 2017). Ultimately, detecting individual profiles of rhythmic abilities in patient populations may play a pivotal role in devising personalized rhythm-based interventions (Dalla Bella, 2020; Dalla Bella et al., 2018). Recent examples show that the individual performance in BPS allows predicting the success of a rhythm-based intervention on motor performance (i.e., gait) in patients with movement disorders (Dalla Bella, Benoit, et al., 2017). Moreover, this information can be integrated in applications capable of providing individualized rhythmic stimulation to patients in real time (Dotov et al., 2019).

## Materials and methods

### Participants

Seventy-nine adults (43 females, 35 right-handed, 4 left-handed and 2 ambidextrous) between 18 and 34 years of age participated in the experiment. Thirty-nine (24 females; mean age = 24.3 years, *SD* = 2.5) were musicians and 40 (19 females; mean age = 23.1, *SD =* 4.0) non-musicians. Musicians were German native speakers recruited via a participant data base at the Max Planck Institute for Human Cognitive and Brain Sciences in Leipzig (Germany), and non-musicians were French speakers recruited in Montpellier (France). Participants had to satisfy the following criteria to be assigned to one of the two groups: 1) they self-classified as musicians or non-musicians, 2) they had practiced during the last year (musicians) or they did not practice during the last year (non-musicians), and 3) did not receive more than 7 years of formal musical training to be considered as non-musicians. A year of formal musical training was defined as a year during which the participant underwent a structured lesson schema of any instrument or voice, either self-taught or by an instructor, and practiced an average of at least 1 hour per week. Musicians reported more years of formal musical training (mean = 7.5 years, *SD* = 2.5) than non-musicians (mean = 1.3 years, *SD* = 2.1; *t*(48.6) = 6.7, *p* < .0001, *d* = 1.9), and more years of informal musical activities (13.1 years, *SD* = 4.4. vs. 3.4 years, *SD =* 4.1; *t*(76.4) = 10.2, *p* < .0001, *d* = 2.3). Informal musical activities consisted in playing one of more musical instrument as an amateur without necessarily having received a formal musical training, which is the case of some non-musicians. The study was approved by ethics committee of the University of Leipzig and by the Euromov Ethics Committee.

### Tests and procedure

We tested participants’ rhythmic abilities by submitting them to the Battery for the Assessment of Auditory and Sensorimotor Timing Abilities (BAASTA; Dalla Bella et al., 2017).

#### Measures of rhythmic abilities

BAASTA consists of a series of 4 perceptual tasks and 5 motor tasks. Perceptual tasks consisted in discriminating single durations (Duration discrimination), detecting deviations from the beat in tone and musical sequences (Anisochrony detection with tones, with music), or in saying whether a superimposed metronome was aligned or not to a musical beat (Beat Alignment Test). Motor tasks involved finger tapping in the absence of stimulation (Unpaced tapping), tapping to the beat of tone and music sequences (Paced tapping with tones, with music), continuing tapping at the pace of a metronome (Synchronization-continuation), and adapting tapping to a tempo change (Adaptive tapping). Participants were tested on all the tasks with a computer version of BAASTA. Auditory stimuli were delivered via headphones (Sennheiser HD201). Task order was fixed (Duration discrimination, Anisochrony detection with tones and music, BAT, for perceptual tasks; Unpaced tapping and Paced tapping to tones and music, for motor tasks). The battery lasted approximately 2 hours.

In the *Duration discrimination* test, two tones (frequency = 1 kHz) were presented successively. The first tone lasted 600 ms (standard duration), while the second lasted between 600 and 1000 ms (comparison duration). Participants judged whether the second tone lasted longer than the first. The goal of the *Anisochrony detection with tones* task was to test the detection of a time shift in an isochronous tone sequence. Sequences of 5 tones (1047 Hz, tone duration = 150 ms) were presented with a mean Inter-Onset Interval (IOI) of 600 ms. Sequences were isochronous (i.e., with a constant IOI) or not (with the 4^th^ tone presented earlier than expected by up to 30% of the IOI). Participants judged whether the sequence was regular or not. The task was repeated using musical stimuli (*Anisochrony detection with music*) that consisted of an excerpt of two bars from Bach’s “Badinerie” orchestral suite for flute (BWV 1067) played with a piano timbre (inter-beat interval = 600 ms). To assess beat perception, the *Beat Alignment Test* (BAT) used 72 stimuli based on 4 regular musical sequences, including 20 beats each (beat = quarter note). Two sequences were fragments from Bach’s “Badinerie”, and 2 from Rossini’s “William Tell Overture”, both played with a piano timbre and played at three different tempos (with 450, 600, and 750-ms Inter-Beat Intervals – IBIs). From the 7^th^ musical beat onward a metronome (i.e., triangle sound) was superimposed onto the music, either aligned or non-aligned to the beat. When non-aligned, the metronome was either phase shifted (with the sounds presented before or after the musical beats by 33% of the music IBI, while keeping the tempo), or period shifted (with the tempo of the metronome changed by ± 10% of the IBI). Participants judged whether the metronome was aligned or not with the musical beat. These perceptual tasks were implemented using Matlab software (version 7.6.0). In the first 3 tasks, there were three blocks of trials and a maximum-likelihood adaptive procedure (MLP) (Green, 1993; MATLAB MLP toolbox, Grassi & Soranzo, 2009) was used to obtain perceptual thresholds (for details, see Dalla Bella et al., 2017). All tasks were preceded by 4 examples and 4 practice trials with feedback. Responses in the perceptual tasks were provided verbally and entered by the Experimenter using the computer keyboard by pressing one of two keys corresponding to a “yes” or “no” response. “Yes” indicated the situation when the participant detected a duration difference, the presence of an anisochrony, or that a metronome was misaligned with the musical beat.

Motor rhythmic abilities were tested with finger tapping tasks. Participants responded by tapping with the index finger of their dominant hand on a MIDI percussion pad. The purpose of the *Unpaced tapping* task was to measure the participants’ preferred tapping rate, and its variability without a pacing stimulus. Participants were asked to tap at their most natural rate for 60 seconds. In two additional unpaced conditions we asked participants to tap as fast as possible for 60 seconds. In the *Paced tapping with tones* task, we asked participants to tap to a metronome sequence, formed by 60 isochronously presented piano tones (frequency = 1319 Hz) at 3 different tempos (600, 450 and 750-ms IOI). Similarly, in the *Paced tapping with music* task participants tapped to the beat of two musical excerpts taken from Bach’s “Badinerie” and Rossini’s “William Tell Overture”. Each musical excerpt contained 64 quarter notes (IBI = 600 ms). Paced tapping trials were repeated twice for each stimulus sequence and were preceded by one practice trial. To test the ability to continue tapping at the rate provided by a metronome, in the *Synchronization-continuation* task participants synchronized with an isochronous sequence of 10 tones (at 600, 450, and 750 ms IOI), and continued tapping at the same rate after the sequence stops, for a duration corresponding to 30 IOIs of the pacing stimulus. The task was repeated twice at each tempo and was preceded by one practice trial. Finally, in the *Adaptive tapping* task, aimed to assess the ability to adapt to a tempo change in a synchronization-continuation task, participants tapped to an isochronous sequence (10 tones). At the end of the sequence (last 4 tones) the tempo either increased, decreased, or remained constant (40% of the trials). The tempo changed by ± 30 or ±75 ms. The task was to tap to the tones in the sequence, to adapt to the tempo change, and to keep tapping at the new tempo after the stimulus stopped for a duration corresponding to 10 IOIs. After each trial, participants judged whether they perceived a change in stimulus tempo (acceleration, deceleration, or no change). The responses were communicated verbally and entered by the Experimenter. Trials were divided into 10 experimental blocks (6 trials x 10 block overall) and presented in random order. A training block preceded the first experimental trial. In all the motor tasks, the performance was recorded via a Roland SPD-6 MIDI percussion pad. Stimulus presentation and response recording was controlled by MAX-MSP software (version 6.0). A MIDI response latency of 133 ms (Leipzig testing) and 100 ms (Montpellier testing) was subtracted from the tapping data before further analysis.

### Analyses

#### BAASTA

For Duration discrimination, Anisochrony detection with tones and with music, we calculated mean thresholds (percentage of the standard duration in the three task) across the three blocks or trials. We rejected the blocks including more than 1/3 of False Alarms, when a difference in duration, or that the sequence beat was irregular, was reported while there was no difference/no deviation from isochrony in the stimulus. In the BAT, we calculated the sensitivity index (*d’*) for the entire set of 72 stimuli, and separately for each or the 3 tempos (medium, fast, and slow). *d’* was calculated based on the number of Hits (when a misaligned metronome was correctly detected) and False alarms (when a misalignment was erroneously reported).

Motor data obtained from tapping tasks were pre-processed as follows (as in Dalla Bella et al., 2017; Sowiński & Dalla Bella, 2013). We discarded taps leading to inter-tap intervals (ITIs) smaller than 100 ms (artifacts) and outlier taps were discarded. An outlier was defined as a tap for which the ITI between the actual tap and the preceding tap was smaller than Q1 – 3*Interquartile range (IQR) or greater than Q3 + 3*IQR, where Q1 is the first quartile and Q3 is the third quartile. We calculated the mean inter-tap interval (ITI, in ms) and motor variability (coefficient of variation of the ITI – CV ITI – namely, the *SD* of the ITI / mean ITI) for Unpaced tapping (spontaneous, slow, and fast), Paced tapping, and for the Synchronization-continuation tasks. Moreover, we analyzed synchronization in the Paced tapping task using circular statistics (Fisher, 1993; Circular Statistics Toolbox for Matlab, Berens, 2009; for use in BAASTA see Dalla Bella et al., 2017). Tap times were coded as unitary vectors with angles relative to the pacing event (tone or musical beat) on a 360° circular scale (corresponding to the inter-stimulus interval). The mean resultant vector *R* was calculated from the unit vectors corresponding to all the taps in a sequence. We used two indexes of synchronization performance: the length of vector *R* (from 0 to 1) and its angle (*θ* or relative phase, in degrees). Vector length indicates whether the taps are systematically occurring before or after the pacing stimuli (*synchronization consistency*); 1 refers to maximum consistency (no variability), and 0, to a random distribution of angles around the circle (i.e., lack of synchronization). The angle of vector *R* (*θ* or relative phase, in degrees) indicates *synchronization accuracy*, namely whether participants tapped before (negative angle) or after (positive angle) the pacing event. Accuracy was calculated only if participants’ synchronization performance was above chance (null hypothesis = random distribution of data points around the circle), as assessed with the Rayleigh test for circular uniformity (Fisher, 1993; Wilkie, 1983). The null hypothesis is rejected when *R* vector length is sufficiently large according to this test. Vector length data was submitted to a logit transformation (e.g., Falk et al., 2015) before conducting further analyses. Additionally, for the Synchronization-continuation task, we calculated measures of central (or timekeeper) variance and motor variance based on Wing-Kristofferson’s model (Wing & Kristofferson, 1973). In both Paced tapping and Synchronization-continuation tasks, the results in the two trials were averaged and submitted to further analyses.

Finally, we analyzed the data from the Adaptive tapping task by first calculating an adaptation index (see Schwartze et al., 2011). To do so, we fitted a regression line to obtain the slope of ITIs functions relative to the final sequence tempo; the value of the slope corresponds to the adaptation index. When the value is 1, the adaptation is perfect; lower and higher values than 1 indicate undercorrection and overcorrection, respectively. We calculated this index separately for tempo acceleration (i.e., faster tempi with final sequence IOIs < 600 ms) and tempo deceleration (slower tempi with final sequence IOIs > 600 ms). In addition, error correction was portioned into the two contributors, phase correction and period correction (Repp & Keller, 2004). Phase and period correction were estimated from the two parameters alpha and beta of the fitted two-process model of error correction (as in Schwartze et al., 2011). Finally, in the same task we calculated the sensitivity index (*d’*) for detecting tempo changes based on the number of Hits (when a tempo acceleration or deceleration was correctly detected) and False Alarms (when a tempo acceleration or deceleration was reported while there was no change or the opposite change). The full list of the 55 variables (7 perceptual, 48 motor) obtained from BAASTA and used as input database for clustering and data modelling is provided in Supplementary materials.

#### Definition of a model of rhythmic abilities (Sparse Learning and Filtering)

The 55 variables from BAASTA collected from a group of 39 musicians and 40 non-musicians served as the starting point to define a model of rhythmic abilities, for classifying the two groups. These variables formed a vector **x** = (*x*_1_,…,*x*_*d*_) in our classification model. Given a set of pairs (*x_k_*, *y*_*k*_),*k* = 1,…,*n*, we looked for a relation *y* = *f*(*x*) that fits the answers *y*_*k*_ by *f*(*x_k_*) ; *y* ; takes value 1 for musicians and 0 for non-musicians. We denote by **Y** the vector of all answers *y*. Finding this relation represents a supervised classification problem as the answers *y*_k_ are known. We addressed this problem using machine learning. One advantage of machine learning is that it does not need *a priori* hypotheses. However, several methods (e.g., neural nets) behave like a “black box” and do not provide any insight about the process and relations leading to successful classification. In this study, to avoid this issue, we looked for an explicit linear relation (see equation 3).

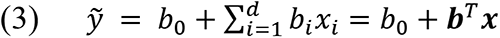

Here,**b** = (*b*_1_,…,*b_d_*)is the vector of regression coefficients and b_0_is the intercept. By **d** we refer to the dimension or number of variables. The symbol **T** represents the transpose of vector b. By 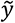 we refer to the approximation of the known response. This is a linear regression model. The classification follows the rule:

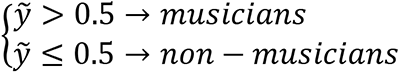

An important issue of all statistical learning models is data dimensions (*d*). In our case, the dimension is large (*d*=55) for a relatively limited sample size (*n*=79). For this reason, one of our goals is to uncover a minimal set of measures capable of classifying musicians and non-musicians, assuming that some variables are redundant and some variables may contribute more than others to classification. To this aim, we employed a sparse learning and filtering (SLF) method (Combettes & Pesquet, 2007; Jiu et al., 2016). This approach presents two advantages: (i) the selected variables are easier to interpret; (ii) for a given number of observations, the statistical power of the prediction increases with the reduction of the dimension (Faul et al., 2009). SLF follows the principle of the lasso method (Tibshirani, 2011) by minimizing both the regression error and the penalized sum of absolute values of coefficients (see equation 4).

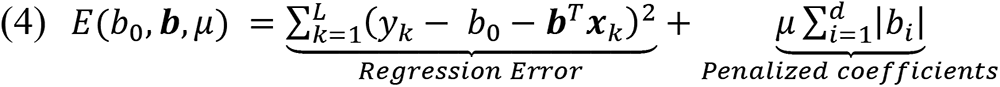

Recent methods of sparse learning optimize a proximal version of this error function which turns to be smooth and allow achieving convergence for well-chosen parameters (Combettes & Pesquet, 2007; Jiu et al., 2016, 2021). Notably, these proximal optimization methods have proven to be successful in supervised classification for big dimension data (d∼10^6^) (Jiu et al., 2016, 2021). is the number of observations that serve to learn the coefficients (b_0_, **b**). Sparse learning methods have the property to bring the coefficients *b*_i_ to 0by tuning the penalizing factor *μ* ≥ 0. The bigger the penalization *μ*, the more *b*_i_ tends to converge to 0. The limit case *μ =* 0 turns to be the basic linear regression and *μ* = ∞ puts all *b*_i_ = 0 thus leading to the constant model 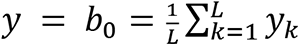 . The optimal *μ* is found by a trade-off between the classification error and the number of non-zero coefficients *b*_i_ .

We used a standard learning protocol (supervised classification) to assess the classification of musicians and non-musicians. We divided the observations into three subsets stratified according to each class (i.e., with balanced percentages of musicians and non-musicians), namely a Train set (60%), a Validation set (20%), and a Test set (20%). The Train set is used to minimize *E*(*b*_0_,**b**,*μ*) (see equation 4) and served to select the variables. The Validation set served as stopping criterion: when prediction errors on Train and Validation sets are approximately equal, the training iterations are stopped. This prevents the model from overfitting the data, another well-known issue of learning methods. The data in the Test set was used to assess the prediction behavior of the model for new data. We used SLF in all the three models: Perc, based on the entire set of perceptual variables (7 out of 55); Motor, based on the entire set of motor variables (48 out of 55); PMI, based on the selected variable as an outcome of models Perc and Motor, also including the interaction between the selected perceptual and motor variables. In Model PMI, we included the interaction between perceptual and motor variables selected for this model, as one of the goals of the study was to assess the relation between perceptual and motor rhythmic abilities in musicians and non-musicians.

Starting from model PMI, we derived profiles of rhythmic abilities for each group, expressed by undirected graphs. The nodes in each graph indicate the variables selected by the model, and the edges represented their interactions. We estimated the contribution of variables and interactions separately to the classification of musicians and non-musicians. To do so we calculated the squared correlation coefficient between each variable or interaction and the prediction and express that as a percentage of the total explained variance in the group. We used this classification procedure based on model PMI to classify musicians and non-musicians, and to classify the two subgroups of non-musicians.

#### Unsupervised learning (modular clustering)

To uncover clusters within the group of non-musicians we considered the participants as a collection of nodes *V*, and link each couple of nodes to form the set of edges *E*. We thus obtained a network that can be easily represented mathematically by a similarity matrix *S*. The value of each entry *S(v,w)* is the similarity between individuals *v* and *w* . If **x** and **y** are the measures for these two individuals then, in our study, their similarity is given by the correlation coefficient *S(v,w)* = *corr*(**x,y**) . We processed these correlations using modularity analysis (Newman, 2006; Rubinov & Sporns, 2010). Modularity quantifies the degree to which a group can be subdivided into clearly delineated and non-overlapping clusters. High modularity reflects strong within-cluster links and weaker links between clusters. We maximized the well-known Newman’s modularity that measures the clustering quality versus a null model. The degree of node *v* is defined by 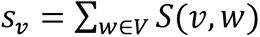 and 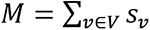 is the sum of degrees. For a given partition *g* of the node’s set *V*, the Modularity (*Q*) is defined as indicated in equation 5.

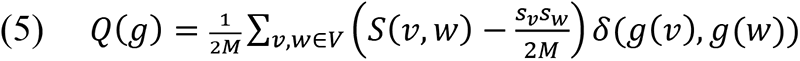

Here, 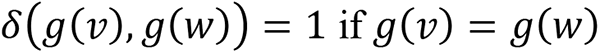 if *g*(*v*) = *g*(*w*) (that is, *v* and *w* are in the same cluster) and 0 otherwise. The null model is given by the value 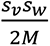 that is the probability for a random edge to appear between *v* and *w*. Then, the clustering algorithm searches for a partition *g* that maximizes *Q*(*g*) (equation 5).

Modularity maximization led to partition the group of non-musicians into two clusters, each including 20 participants (for a representation of heat matrix showing that intra-cluster similarities are larger than inter-cluster similarities in the dataset, see Supplementary materials).

## Notes

This research was funded by the European Community’s Seventh Framework Programme (EBRAMUS project, FP7 Initial Training Network, grant agreement no. 238157) and by FEDER funds (Languedoc-Roussillon Region, BAASTA-FEDER project) to SDB and SAK; by Canada Research Chair program funding to SDB (CRC in music auditory-motor skill learning and new technologies), and by international mobility funds from IMT Mines Alès to SJ.

