## Supplementary materials for "Unravelling individual rhythmic abilities using machine learning"

### 1) List of variables

Full list of variables entered initially in the analysis.

| Var # | Variable name | Task | Task category | Outcome measure |
|---|---|---|---|---|
| 1 | Dur_discrim_Thresh | Duration discrimination | Perceptual | Threshold for duration discrimination |
| 2 | Anisoc_det_tones_Thresh | Anisochrony detection (tones) | Perceptual | Threshold for anisochrony detection with tones |
| 3 | Anisoc_det_music_Thresh | Anisochrony detection (music) | Perceptual | Threshold for anisochrony detection with music |
| 4 | BAT_all_Dprime | Beat Alignment Test (BAT) | Perceptual | *d'* (sensitivity index) based on all trials (72) |
| 5 | BAT_fast_Dprime | Beat Alignment Test (BAT) | Perceptual | *d'* (sensitivity index) based on trials at fast tempo (24) |
| 6 | BAT_med_Dprime | Beat Alignment Test (BAT) | Perceptual | *d'* (sensitivity index) based on trials at medium tempo (24) |
| 7 | BAT_slow_Dprime | Beat Alignment Test (BAT) | Perceptual | *d'* (sensitivity index) based on trials at slow tempo |
| 8 | Unpaced_spont_ITI | Unpaced tapping | Motor | Rate (inter-tap interval) of spontaneous tapping |
| 9 | Unpaced_spont_CV | Unpaced tapping | Motor | Variability (coefficient of variation of the inter-tap interval) of spontaneous tapping |
| 10 | Unpaced_slow_ITI | Unpaced tapping | Motor | Rate of slow tapping |
| 11 | Unpaced_slow_CV | Unpaced tapping | Motor | Variability of slow tapping |
| 12 | Unpaced_fast_ITI | Unpaced tapping | Motor | Rate of fast tapping |
| 13 | Unpaced_fast_CV | Unpaced tapping | Motor | Variability of fast tapping |
| 14 | Paced_metro_450_ITI | Paced tapping (metronome) | Motor | Rate of paced tapping (IOI = 450ms) |
| 15 | Paced_metro_450_CV | Paced tapping (metronome) | Motor | Variability of paced tapping (IOI = 450ms) |
| 16 | Paced_metro_450_Vector_dir | Paced tapping (metronome) | Motor | Accuracy (vector direction) of paced tapping (IOI = 450ms) |
| 17 | Paced_metro_450_Vector_len | Paced tapping (metronome) | Motor | Consistency (vector length) of paced tapping (IOI = 450ms) |
| 18 | Paced_metro_600_ITI | Paced tapping (metronome) | Motor | Rate of paced tapping (IOI = 600ms) |
| 19 | Paced_metro_600_CV | Paced tapping (metronome) | Motor | Variability of paced tapping (IOI = 600ms) |
| 20 | Paced_metro_600_Vector_dir | Paced tapping (metronome) | Motor | Accuracy (vector direction) of paced tapping (IOI = 600ms) |
| 21 | Paced_metro_600_Vector_len | Paced tapping (metronome) | Motor | Consistency (vector length) of paced tapping (IOI = 600ms) |
| 22 | Paced_metro_750_ITI | Paced tapping (metronome) | Motor | Rate of paced tapping (IOI = 750ms) |
| 23 | Paced_metro_750_CV | Paced tapping (metronome) | Motor | Variability of paced tapping (IOI = 750ms) |

| | | | | |
|---|---|---|---|---|
| 24 | Paced_metro_750_Vector_dir | Paced tapping (metronome) | Motor | Accuracy (vector direction) of paced tapping (IOI = 750ms) |
| 25 | Paced_metro_750_Vector_len | Paced tapping (metronome) | Motor | Consistency (vector length) of paced tapping (IOI = 750ms) |
| 26 | Paced_music_badine_ITI | Paced tapping (music) | Motor | Rate of paced tapping (music 1) |
| 27 | Paced_music_badine_CV | Paced tapping (music) | Motor | Variability of paced tapping (music 1) |
| 28 | Paced_music_badine_Vector_dir | Paced tapping (music) | Motor | Accuracy (vector direction) of paced tapping (music 1) |
| 29 | Paced_music_badine_Vector_len | Paced tapping (music) | Motor | Consistency (vector length) of paced tapping (music 1) |
| 30 | Paced_music_ross_ITI | Paced tapping (music) | Motor | Rate of paced tapping (music 2) |
| 31 | Paced_music_ross_CV | Paced tapping (music) | Motor | Variability of paced tapping (music 2) |
| 32 | Paced_music_ross_Vector_dir | Paced tapping (music) | Motor | Accuracy (vector direction) of paced tapping (music 2) |
| 33 | Paced_music_ross_Vector_len | Paced tapping (music) | Motor | Consistency (vector length) of paced tapping (music 2) |
| 34 | Sync_Cont_ 450_ITI | Synchronization-continuation | Motor | Rate of tapping in the continuation phase (IOI = 450ms) |
| 35 | Sync_Cont_450_ CV | Synchronization-continuation | Motor | Variability of tapping in the continuation phase (IOI = 450ms) |
| 36 | Sync_Cont_450_Motor_variance | Synchronization-continuation | Motor | Motor variance, calculated from tapping in the continuation phase (IOI = 450ms) |
| 37 | Sync_Cont_450_Central_variance | Synchronization-continuation | Motor | Central variance, calculated from tapping in the continuation phase (IOI = 450ms) |
| 38 | Sync_Cont_600_ITI | Synchronization-continuation | Motor | Rate of tapping in the continuation phase (IOI = 600ms) |
| 39 | Sync_Cont_600_ CV | Synchronization-continuation | Motor | Variability of tapping in the continuation phase (IOI = 600ms) |
| 40 | Sync_Cont_600_Motor_variance | Synchronization-continuation | Motor | Motor variance, calculated from tapping in the continuation phase (IOI = 600ms) |
| 41 | Sync_Cont_600_Central_variance | Synchronization-continuation | Motor | Central variance, calculated from tapping in the continuation phase (IOI = 600ms) |
| 42 | Sync_Cont_ 750_ITI | Synchronization-continuation | Motor | Rate of tapping in the continuation phase (IOI = 750ms) |
| 43 | Sync_Cont_750_ CV | Synchronization-continuation | Motor | Variability of tapping in the continuation phase (IOI = 750ms) |

| | | | | |
|---|---|---|---|---|
| 44 | Sync_Cont_750_Motor_variance | Synchronization-continuation | Motor | Motor variance, calculated from tapping in the continuation phase (IOI = 750ms) |
| 45 | Sync_Cont_750_Central_variance | Synchronization-continuation | Motor | Central variance, calculated from tapping in the continuation phase (IOI = 750ms) |
| 46 | Adaptive_Adapt_index_accel | Adaptive tapping | Motor | Adaptation index (acceleration trials) |
| 47 | Adaptive_Adapt_index_decel | Adaptive tapping | Motor | Adaptation index (deceleration trials) |
| 48 | Adaptive_Alpha_accel | Adaptive tapping | Motor | Phase correction (alpha coefficient), acceleration trials |
| 49 | Adaptive_Alpha_decel | Adaptive tapping | Motor | Phase correction (alpha), deceleration trials |
| 50 | Adaptive_Beta_accel | Adaptive tapping | Motor | Period correction (beta coefficient), acceleration trials |
| 51 | Adaptive_Beta_decel | Adaptive tapping | Motor | Period correction (alpha), deceleration trials |
| 52 | Adaptive_Dprime_min30 | Adaptive tapping | Motor | *d'* when perceiving a tempo acceleration (IOI - 30ms) |
| 53 | Adaptive_Dprime_min75 | Adaptive tapping | Motor | *d'* when perceiving a tempo acceleration (IOI - 75ms) |
| 54 | Adaptive_Dprime_plus30 | Adaptive tapping | Motor | *d'* when perceiving a tempo deceleration (IOI + 30ms) |
| 55 | Adaptive_Dprime_plus75 | Adaptive tapping | Motor | *d'* when perceiving a tempo deceleration (IOI + 75ms) |

Total number of variables: 55 (7 from perceptual tasks, 48 from motor tasks).

### 2) Classification of musicians (MUS) and non-musicians (n-MUS)

The confusion matrices for the Train, Validation, and Test sets for the three models (Perc, Motor, PMI) are presented in Table 2a, b, and c respectively.

*Table 1a. Confusion matrices for classification based on model Perc.*

| | Train | | Validation | | Test | |
|---|---|---|---|---|---|---|
| | Pred n-MUS | Pred MUS | Pred n-MUS | Pred MUS | Pred n-MUS | Pred MUS |
| Actual n-MUS | 13 | 8 | 8 | 2 | 6 | 3 |
| Actual MUS | 5 | 19 | 0 | 7 | 1 | 7 |

*Table 2b. Confusion matrices for classification based on model Motor.*

| | Train | | Validation | | Test | |
|---|---|---|---|---|---|---|
| | Pred n-MUS | Pred MUS | Pred n-MUS | Pred MUS | Pred n-MUS | Pred MUS |
| Actual n-MUS | 15 | 6 | 7 | 3 | 8 | 1 |
| Actual MUS | 5 | 19 | 0 | 7 | 3 | 5 |

*Table 3c. Confusion matrices for classification based on Model PMI.*

| | Train | | Validation | | Test | |
|---|---|---|---|---|---|---|
| | Pred n-MUS | Pred MUS | Pred n-MUS | Pred MUS | Pred n-MUS | Pred MUS |
| Actual n-MUS | 15 | 6 | 8 | 2 | 8 | 2 |
| Actual MUS | 3 | 21 | 2 | 6 | 0 | 7 |

#### 3) Unsupervised learning (modular clustering) for non-musicians

The result of modularity maximization is a partition of non-musicians into two clusters each including 20 participants. The modularity value is $Q = 0.14$. The following Figure  shows the heat matrix $S$ with participants in their original order and the same matrix with rows and columns permuted to fit the clusters found above. It is easily seen that intra similarities are stronger (brighter) than inter similarities.

The following Figure shows the heat matrix $S$ with participants in their original order and the same matrix with rows and columns permuted to fit the clusters found above. It can be easily seen that intra-cluster similarities are larger (brighter) than inter-cluster similarities.

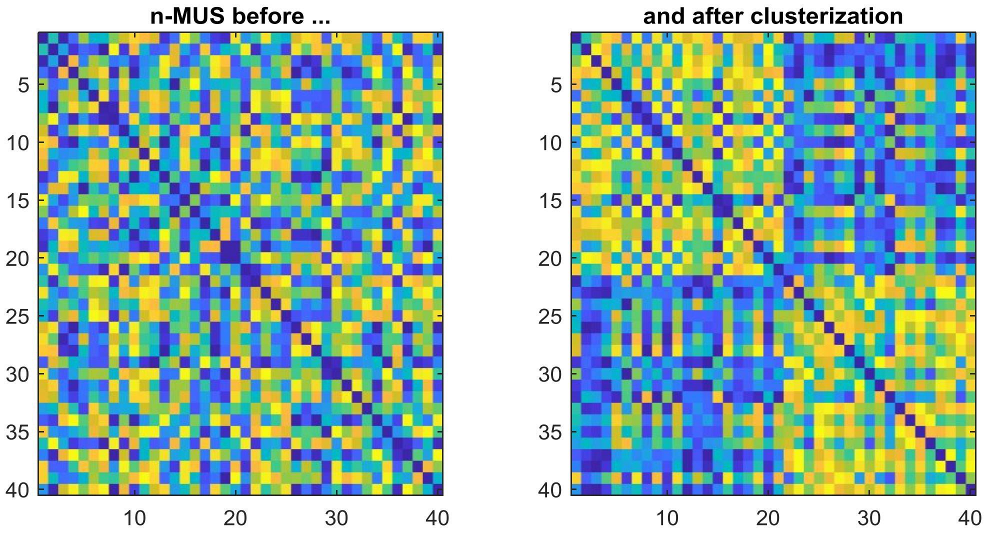

*Figure 1. Heat matrices. The brighter the colour the more similar are individuals. Left: Similarity matrix before clustering; Right: the same matrix with rows and columns reordered by clusters C1 and C2.*

#### 4) Classification of the two subgroups of non-musicians (nMUS-1, nMUS-2)

*Table 3*. Confusion matrix for classification based on Model PMI.

| | Train | | Validation | | Test | |
|---|---|---|---|---|---|---|
| | Pred nMUS-1 | Pred nMUS-2 | Pred nMUS-1 | Pred nMUS-2 | Pred nMUS-1 | Pred nMUS-2 |
| Actual nMUS-1 | 9 | 0 | 5 | 1 | 5 | 0 |
| Actual nMUS-2 | 0 | 12 | 0 | 3 | 0 | 5 |
